# The RhoGAP SPV-1 regulates calcium signaling to control the contractility of the *C. elegans* spermatheca during embryo transits

**DOI:** 10.1101/441386

**Authors:** Jeff Bouffard, Alyssa D. Cecchetelli, Coleman Clifford, Kriti Sethi, Ronen Zaidel-Bar, Erin J. Cram

**Affiliations:** Department of Bioengineering, Northeastern University, Boston, MA; Department of Biology, Northeastern University, Boston, MA; Mechanobiology Institute, National University of Singapore, Singapore; Department of Cell and Developmental Biology, Sackler Faculty of Medicine, Tel-Aviv University, Israel

## Abstract

Contractility of the non-muscle and smooth muscle cells that comprise biological tubing is regulated by the Rho-ROCK and calcium signaling pathways. Although many molecular details about these signaling pathways are known, less is known about how they are coordinated spatiotemporally in biological tubes. The spermatheca of the *C. elegans* reproductive system enables study of the signaling pathways regulating actomyosin contractility in live adult animals. The RhoGAP SPV-1 was previously identified as a negative regulator of RHO-1/Rho and spermathecal contractility. Here, we uncover a role for SPV-1 as a key regulator of calcium signaling. *spv-1* mutants expressing the calcium indicator GCaMP in the spermatheca exhibit premature calcium release, elevated calcium levels, and disrupted spatial regulation of calcium signaling during spermathecal contraction. Although RHO-1 is required for spermathecal contractility, RHO-1 does not play a significant role in regulating calcium. In contrast, activation of CDC-42 recapitulates many aspects of *spv-1* mutant calcium signaling. Depletion of *cdc-42* by RNAi does not suppress the premature or elevated calcium signal seen in *spv-1* mutants, suggesting other targets remain to be identified. Our results suggest SPV-1 works through both the Rho-ROCK and calcium signaling pathways to coordinate cellular contractility.

**Highlight Summary:** Through *in vivo* imaging of the calcium sensor GCaMP, we show that the RhoGAP SPV-1 is a key regulator of calcium signaling in the *C. elegans* spermatheca. Our data suggests SPV-1 acts at least partially through the small GTPase CDC-42 to modulate calcium signaling, while also acting on RHO-1 to modulate Rho-ROCK signaling. This places SPV-1 as a central regulator of cellular contractility.

## Introduction

Animals are full of biological tubing, including blood and lymphatic vessels, lung airways, salivary glands, digestive canals, and urinary and reproductive tracts. Actomyosin contractility plays a central role in the functioning of these biological tubes, which must dilate and contract with the proper timing and magnitude to generate appropriate responses to changing biological states (Reviewed in Sethi et al., 2017). Consequences of misregulated biological tubes can be seen in conditions such as heart disease, hypertension, and asthma (Lavoie et al., 2009; Seguchi et al., 2007; Uehata et al., 1997; Wettschureck and Offermanns, 2002). Previous research has focused on how biochemical factors such as acetylcholine, serotonin, and nitric oxide regulate tissue function in biological tubes, and elucidated many of the downstream signaling mechanisms that regulate cellular and actomyosin contractility. However, mechanical factors such as flow and pressure are also known to regulate tissue function in biological tubes (Gunst et al., 2003; Smiesko and Johnson, 1993; Smith et al., 2003), and the mechanisms by which these mechanical signals regulate cellular contractility are not as well understood. *In vivo* studies suggest that mechanical cues play a critical role in regulating and coordinating actomyosin contractility (Munjal and Lecuit, 2014).

The *C. elegans* spermatheca (Figure 1A) is a powerful *in vivo* model system for the study of how cells regulate actomyosin contractility and produce coordinated tissue-level responses to mechanical input. The spermatheca is part of the hermaphrodite gonad, a simple, tubular organ consisting of the oviduct (containing germ cells and oocytes and enclosed by contractile sheath cells), the spermatheca, and the uterus (Kimble and Hirsh, 1979). The spermatheca, consisting of a single layer of 24 myoepithelial cells (Figure 1B), is the site of sperm storage and fertilization. Sheath cell contractions propel the oocyte into the spermatheca, dramatically stretching the cells of the distal neck and spermathecal bag and initiating a process that culminates in spermathecal cell contraction, spermatheca uterine (sp-ut) valve dilation, and expulsion of the fertilized egg into the uterus. This ovulation cycle, including oocyte entry, transit through the spermatheca, and expulsion into the uterus, repeats roughly every 20 minutes until ~150 eggs have been produced by each gonad arm. Because the nematode is transparent, and the cells of the spermatheca are clearly visible, the entire ovulatory process can be visualized in live animals using time-lapse microscopy.

**Figure 1.**
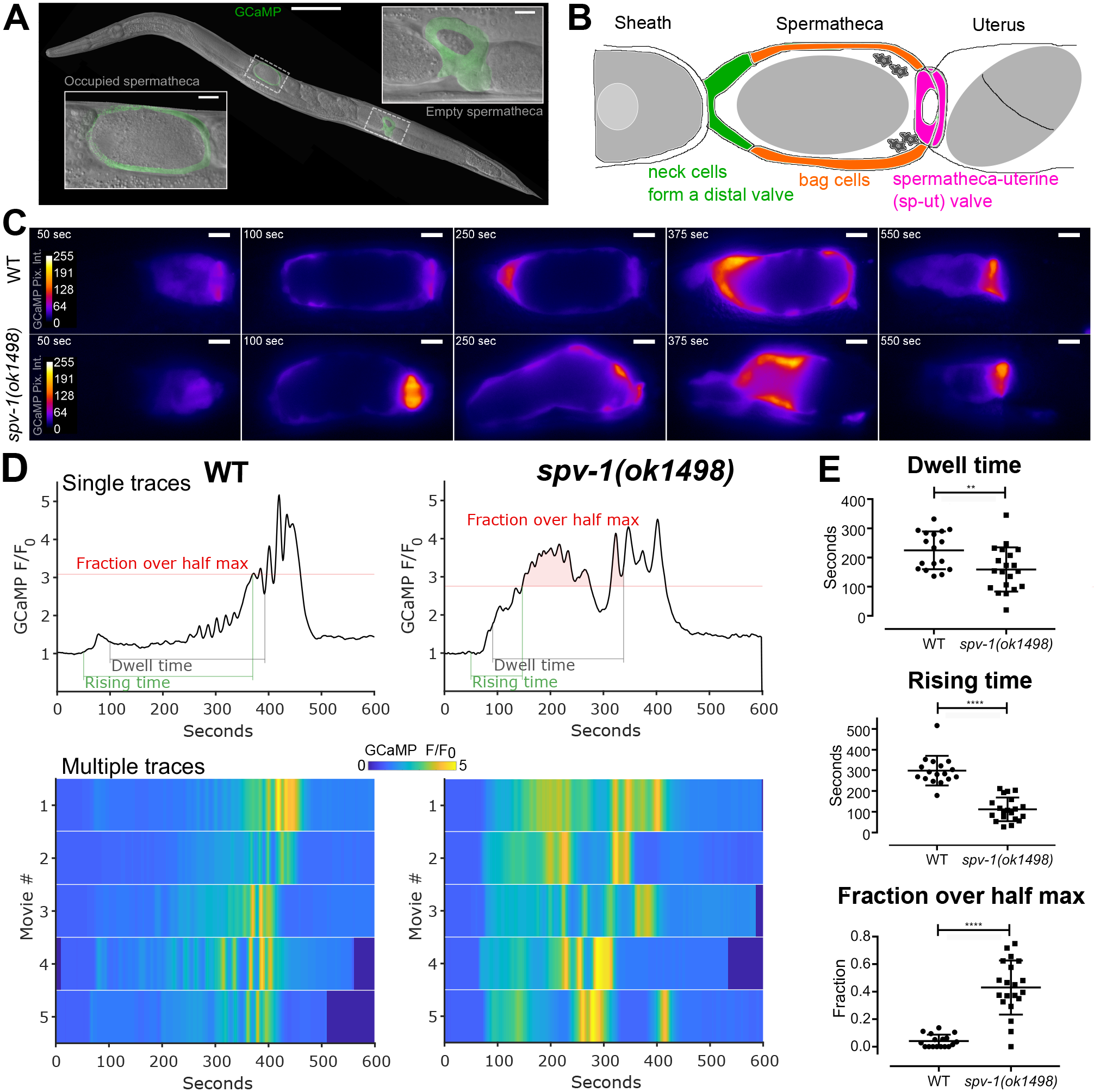
Loss of SPV-1 alters calcium signaling in the spermatheca. (A) An adult nematode with labeled spermathecae, showing one with an embryo inside (left), and one without an embryo inside (right). The scale bar in the large image is 100 μm, scale bars in the insets are 10 μm. (B) Tissue-level schematic cartoon of the spermatheca. The spermatheca consists of 3 distinct regions: the cells closest to the sheath form a neck, or distal valve, that constricts to enclose the newly entered embryo, the central cells form a bag that accomodates the embryo during fertilization and egg shell deposition, and the spermatheca and uterus are connected by the sp-ut valve. (C) Still frames from GCaMP movies of embryo transits in wild-type (top) and *spv-1(ok1498)* (bottom) animals. Movies were temporally aligned to the start of oocyte entry at 50 seconds. (D) GCaMP time series generated from GCaMP movies (Movie 1), with metrics highlighted. Dwell time is a tissue function metric calculated as the time from the closing of the distal valve to the opening of the sp-ut valve. Rising time is a calcium signaling metric measuring the time from the opening of the distal valve to the first time point where the time series reaches half its maximum, and fraction over half max is a calcium signaling metric measuring how much of the dwell time is spent over the half maximum. Lower panels show heat maps of 5 time series for each condition. The top line of the heat map is the time series in the upper panel. (E) Quantification of metrics from time series. Error bars display standard deviation, p-values were calculated using Welch’s t-test: **p < 0.01, ****p < 0.0001.

Contraction of the spermatheca is driven by the well conserved Rho-ROCK and calcium signaling pathways, similar to those observed in non-muscle and smooth muscle cells (Brozovich et al., 2016; Pelaia et al., 2008; Sethi et al., 2017). These two pathways act in concert to regulate the levels of phosphorylated myosin (Somlyo and Somlyo, 2003). The calcium signaling pathway requires activated phospholipase C (PLC-1), which cleaves the membrane lipid phospatidyl inositol bisphosphate (PIP_2_) to generates 1,4,5 triphosphate (IP_3_) and diacylglycerol (DAG). IP_3_ stimulates the release of calcium from the endoplasmic reticulum (ER) by binding to the IP_3_ receptor (ITR-1), and this elevated cytosolic calcium activates myosin light chain kinase (MLCK-1), which phosphorylates the regulatory light chain of myosin (MLC-4). The Rho-ROCK pathway acts via RhoA (RHO-1), with active, GTP-bound Rho activating Rho kinase/ROCK (LET-502), which phosphorylates and activates myosin and also inhibits myosin phosphatase (MEL-11), resulting in increased phosphorylation and activation of non-muscle myosin II (NMY-1), and contraction of actomyosin fibers. Activation and coordination of both the Rho-ROCK and calcium signaling pathways is required for successful transits of embryos through the spermatheca (Kovacevic et al., 2013; Tan and Zaidel-Bar, 2015). Despite our detailed molecular understanding of these two central pathways regulating contractility, little is known about how these two pathways are integrated to regulate actomyosin contractility at the cellular level.

We have developed the *C. elegans* spermatheca as a model for understanding how cells within a tissue coordinately regulate actomyosin contractility (Kelley et al., 2018; Kovacevic et al., 2013; Tan and Zaidel-Bar, 2015; Wirshing and Cram, 2017). Using the genetically encoded calcium sensor, GCaMP, we have shown that the stretch of incoming oocytes triggers a flash of calcium in the sp-ut valve, followed by dynamic oscillations across the central bag which culminate in contraction and expulsion of the fertilized embryo through the sp-ut valve and into the uterus (Kovacevic et al., 2013). Phospholipid signaling, communication through gap junctions, and the mechanosensor FLN-1/filamin (Kovacevic and Cram, 2010; Kovacevic et al., 2013) are all required for proper spatiotemporal regulation of calcium signaling. Spatiotemporal control of Rho-ROCK signaling is also required for coordinated regulation of contractility in the spermatheca. We have shown that the RhoGAP SPV-1 controls the spatiotemporal activation of RHO-1, and regulates tissue contractility through LET-502/ROCK (Tan and Zaidel-Bar, 2015). Disruption of these spatiotemporal regulators often results in damage to embryos, reflux of embryos back into the oviduct, or embryos that are trapped in the spermatheca and unable to exit.

In this study, we demonstrate that the RhoGAP SPV-1 is necessary for proper calcium signaling in the *C. elegans* spermatheca. We further find that SPV-1 is a major regulator of spatiotemporal aspects of spermathecal calcium signaling, controlling the timing and magnitude of calcium signaling in the spermathecal bag and sp-ut cells. Expression of constitutively active alleles of RHO-1 and CDC-42 recapitulate the contractility and calcium release phenotypes, respectively, of loss of *spv-1*. This places SPV-1 as a central regulator of both major signaling pathways that modulate actomyosin contractility.

## Results

### SPV-1 regulates calcium signaling in the spermatheca during embryo transits

To determine if SPV-1 plays a role in spermathecal calcium signaling we used a mutant allele of *spv-1, ok1498*, which causes a 577-bp frameshift deletion in the RhoGAP domain, resulting in loss of function of the SPV-1 protein (Tan and Zaidel-Bar, 2015). We imaged embryo transits in wildtype and *sp-1(ok1498)* nematodes expressing the genetically encoded calcium sensor GCaMP3 in the spermatheca (Figure 1C; Movie 1). To characterize the *spv-1* mutant calcium signaling phenotype, we generated GCaMP time series from the embryo transit movies by calculating the average pixel intensity of each frame (Figure 1, C and D; Movie 1). Plotting time series from many animals as heat maps reveals that *spv-1(ok1498)* embryo transits consistently show a rapid increase in the onset of calcium activity at the start and elevated calcium levels throughout (Figure 1D).

To quantify these differences and enable statistical analysis over many embryo transits, we identified one tissue function metric from the movies, dwell time, and two calcium signaling metrics extracted from the GCaMP time series, rising time and fraction over half max (Figure 1D; Movie 1). The tissue function metric, dwell time, is defined as the time from distal neck closure to sp-ut valve opening. Dwell time measures the amount of time the embryo remains enclosed by the spermatheca. In *spv-1* mutants, faster transit of the embryo through the spermatheca results in a reduced dwell time (Figure 1E). This result is in agreement with a previous observation that embryo transits are faster when SPV-1 is lost (Tan and Zaidel-Bar, 2015). The first calcium signaling metric, rising time, is defined as the time from the start of oocyte entry to the first time point where the GCaMP time series crosses the half maximum. Rising time captures the rate of increase in calcium signal at the start of embryo transit. In *spv-1* mutants, a rapid rise in calcium is observed immediately upon entry (Figure 1, D and E). The second calcium signaling metric, fraction over half max, is defined as the duration of the dwell time over the GCaMP half maximum value divided by the total dwell time. This metric captures the level of calcium throughout embryo transits. In *spv-1* mutants, fraction over half max is higher than in wild type animals (Figure 1, D and E). Taken together, these metrics suggest SPV-1 regulates calcium activity in the spermatheca by dampening calcium signaling, keeping cytosolic calcium at low levels until late in embryo transits.

### SPV-1 overexpression results in low calcium signaling and embryo trapping

To further understand how SPV-1 regulates calcium signaling, we expressed an SPV-1::mApple translational fusion in the *spv-1* mutant background. Animals expressing both the mApple fluorophore and GCaMP were imaged during embryo transits (Figure 2, A and B). High levels of SPV-1::mApple fluorescence intensity were associated with failure of embryos to exit the spermatheca (referred to here as embryo trapping) and low magnitude calcium activity (Figure 2, C and D), suggesting that overexpression of SPV-1 results in disrupted tissue function and calcium signaling phenotypes. We next crossed *spv-1::mApple* into the wildtype background, and observed embryo trapping and low calcium signaling in 100% of movies recorded (n=5, data not shown). In control experiments, similar levels of mApple intensity did not coincide with embryo trapping (Figure 2, E and F). These data suggest that high levels of SPV-1 inhibit calcium release and spermathecal tissue contractility.

**Figure 2.**
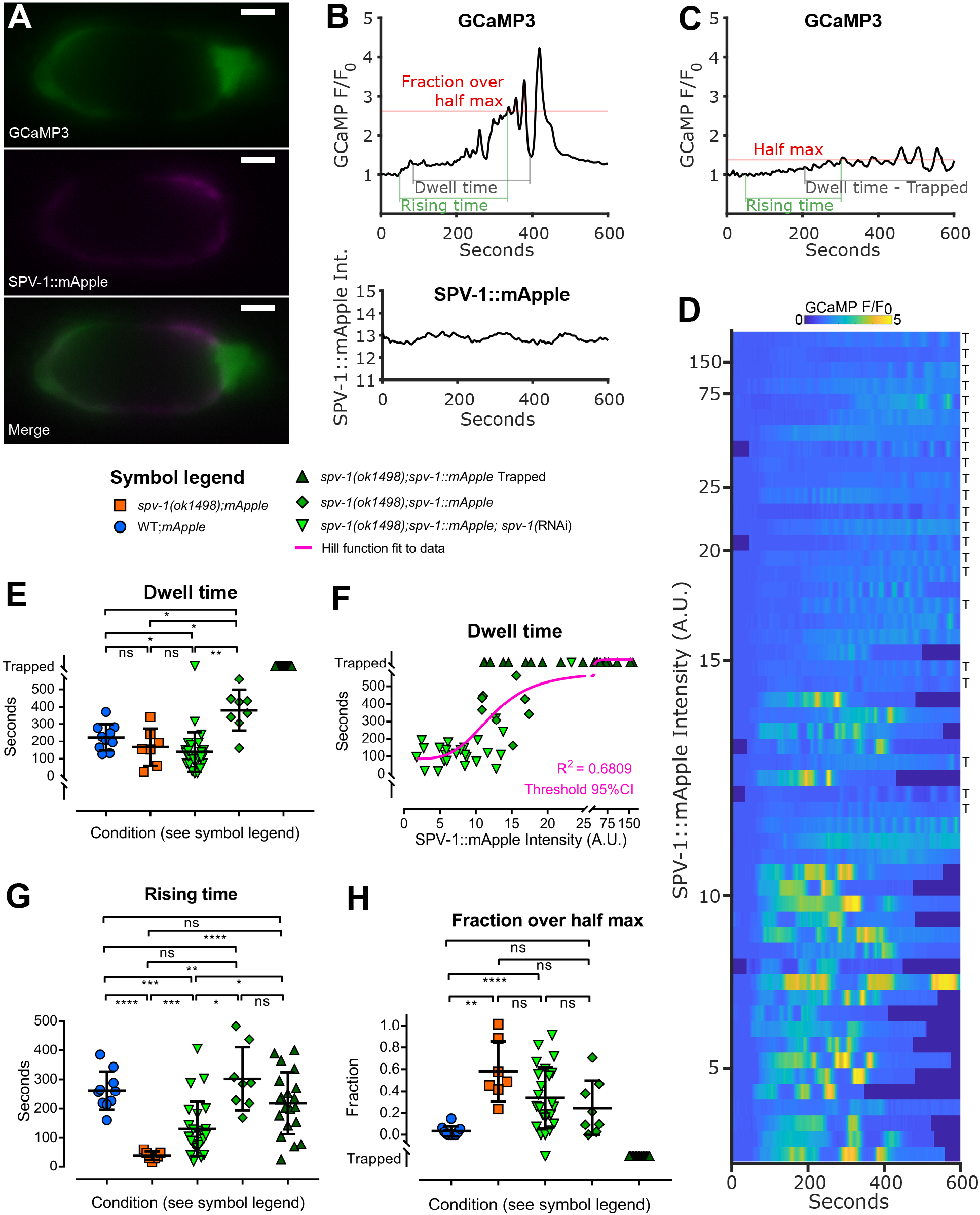
Spermathecal tissue function and calcium signaling exhibit a threshold response to SPV-1::mApple. (A) Still images from a dual-labeled spermatheca expressing GCaMP and SPV-1::mApple. Scale bars are 10 μm (B) GCaMP time series of normalized average pixel intensity, F/ F_0_ (top), and SPV1::mApple time series of raw average pixel intensity (bottom), from a single embryo transit movie over the same spatial frame and time. (C) A representative GCaMP time series from an SPV-1::mApple expressing spermatheca containing a trapped embryo, corresponding to the top row of the heat map in panel D. (D) A heat map showing GCaMP time series from 53 embryo transit movies of varying SPV-1::mApple intensity, with the highest SPV-1::mApple intensity at the top and decreasing with each row. Rows labeled with ‘T’ indicate trapped. (E) Dwell times plotted as a function of condition. (F) Dwell times plotted as a function of SPV-1::mApple intensity. The threshold value from the fitted Hill function is 12.2, with the 95% confidence interval from 10.1 to 14.8. (G) Rising times plotted as a function of condition. (H) Fractions over half max plotted as a function of condition. In panels E, G, and H error bars display standard deviation, p-values were calculated using Welch’s ANOVA with Games-Howell multiple comparison: ns, p ≥ 0.05, *p < 0.05, **p < 0.01, ***p <0.001, ****p < 0.0001.

### Spermathecal tissue function and calcium signaling exhibit a threshold response to SPV-1::mApple levels

To examine how spermathecal tissue function and calcium signaling respond to varying amounts of SPV-1, we used diluted RNAi against *spv-1* to reduce SPV-1::mApple levels. These treatments generated *spv-1(ok1498); spv-1::mApple* animals with a range of SPV-1::mApple levels, measured by mApple fluorescence intensity. Spermathecal calcium signaling and embryo transit timing exhibit a threshold response to SPV-1::mApple levels, in which low levels of SPV-1::mApple result in short dwell times and *spv-1* mutant-like calcium signaling with a rapid onset of elevated calcium activity, whereas high levels of SPV-1::mApple induce the overexpression phenotypes with embryo trapping and low calcium activity (Figure 2, D and E). Plotting dwell times as a function of SPV-1::mApple intensity clearly displays this threshold behavior (Figure 2F), which can be modeled using a Hill function (A.V. Hill, 1910) to determine where, and how rapidly, a system switches from one state to another (Monod et al., 1965). To explore this we fit a Hill equation to these dwell time data points (Figure 2F), and extracted a threshold value of 12.2 for SPV-1::mApple intensity, with a 95% confidence interval from 10.1 to 14.8. This window of SPV-1::mApple intensity values coincides with the switch from mutant to overexpression behavior in both tissue function (Figure 2, E and F) and calcium signaling (Figure 2, D, G and H).

### SPV-1 regulates spatiotemporal aspects of calcium signaling

In wildtype embryo transits, calcium signaling starts with a single pulse in the sp-ut valve, followed by a quiet period after the oocyte enters, then increasing pulses are initiated in the distal neck and travel across the spermathecal bag, culminating in intense calcium pulses that activate contractions that expel the embryo into the uterus (Kovacevic et al., 2013). In contrast, in *spv-1(ok1498)* embryo transits calcium rises immediately upon oocyte entry, lacks spatial organization, and is elevated compared to wildtype signal, sporadically crashing to a low level and rising again multiple times before the embryo is expelled (Figure 1, B and C; Figure 3A; Movie 1).

**Figure 3.**
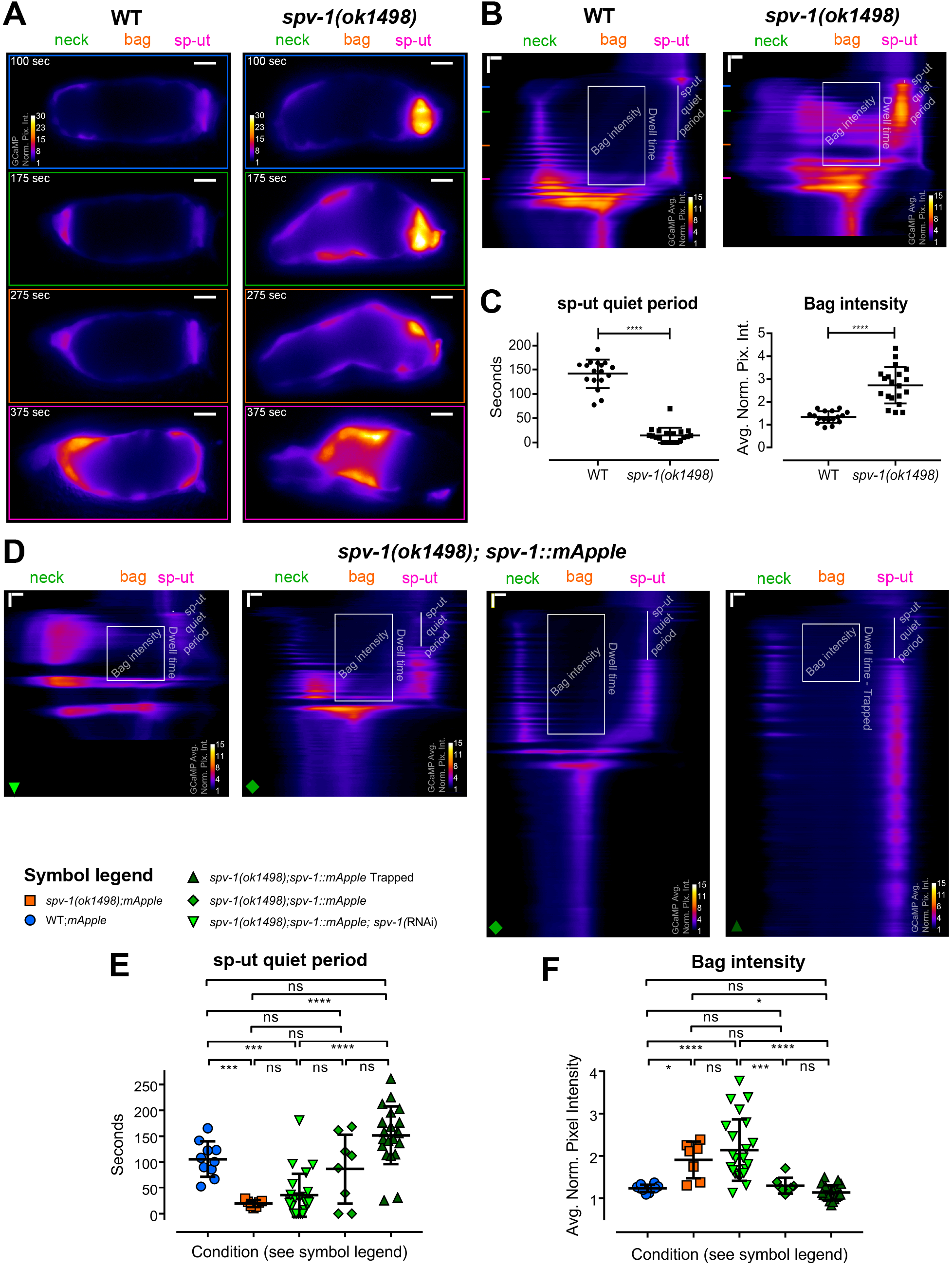
SPV-1 regulates spatiotemporal aspects of calcium signaling. (A) Individual frames from wildtype (left) and *spv-1(ok1498)* (right) embryo transit movies. All frames follow the color scale indicated in the top frame, scale bars are 10 μm. (B) Kymograms of the movies in panel A, generated by averaging over the columns of each movie frame, display the variation in average calcium signaling from the distal valve on the left to the sp-ut valve on the right, with time progressing down (Movie 2). Horizontal scale bars are 5 μm, vertical scale bars are 50 seconds. Colored lines on the left side of the kymograms correspond to the individual frames in panel A, annotations show the two spatiotemporal calcium signaling metrics used for analysis. The sp-ut quiet period measures the low calcium signaling of the sp-ut valve after oocyte entry, which is lost in *spv-1* mutants. Bag intensity measures the average normalized fluorescence intensity of a 25 μm wide region in the bag section of the spermatheca during the dwell time. (C) Quantification of metrics. Error bars display standard deviation, p-values were calculated with Welch’s t-test: ****p < 0.0001. (D) Kymograms from embryo transit movies with SPV-1::mApple intensities of 2.5, 12.6, 16.9, and 78.5, from left to right. Horizontal scale bars are 5 μm, vertical scale bars are 50 seconds. (E) sp-ut quiet periods plotted as a function of condition. (F) Bag intensities plotted as a function of condition. In panels E and F error bars display standard deviation, p-values were calculated using Welch’s ANOVA with Games-Howell multiple comparison: ns, p ≥ 0.05, *p < 0.05, ***p <0.001, ****p < 0.0001.

To visualize and analyze these spatiotemporal changes we generated kymograms in which each frame of the movie is condensed into a horizontal line, along the distal to proximal axis of the spermatheca, by averaging over the columns of that frame. Lines representing the individual frames are then stacked vertically, generating kymograms that display the spatial variation of the GCaMP signal from the distal neck to the sp-ut valve over the entire movie (Figure 3, A and B; Movie 2). Wildtype kymograms indicate calcium activity is restricted to the distal (neck) and proximal (sp-ut) ends of the spermatheca during the dwell time. In contrast, *spv-1(ok1498)* kymograms depict increased activity in the middle of the spermathecal bag as well as in the sp-ut valve (Figure 3B).

To quantify these differences, we identified two prominent spatiotemporal calcium signaling metrics that are consistently altered in *spv-1* mutant kymograms (Figure 3, B and C). The first is the sp-ut quiet period. In wildtype kymograms, the sp-ut valve displays a single calcium transient upon oocyte entry, followed by an extended period of low calcium. In *spv-1(ok1498)* kymograms, the sp-ut valve displays an immediate rise in calcium followed by sustained calcium. The sp-ut quiet period metric quantifies this difference by measuring the time from the end of the initial calcium transient to the next rise in sp-ut calcium. The second spatiotemporal calcium signaling metric is bag intensity. In wildtype kymograms, the central spermathecal cells exhibit low calcium activity that increases and pulses as the tissue expels the egg. In *spv-1(ok1498)* kymograms these central cells exhibit rapid high calcium levels that stay elevated while the spermatheca is occupied. The bag intensity metric quantifies this difference by measuring the average pixel intensity of a 25 μm region of the spermathecal bag over the dwell time. Both spatiotemporal metrics differ significantly between wildtype and *spv-1(ok1498)* animals (Figure 3C).

We next examined kymograms from the movies with varying SPV-1::mApple intensities (Figure 3D). As expected, low levels of SPV-1::mApple result in short sp-ut quiet periods and high bag intensities, whereas high levels of SPV-1::mApple result in long sp-ut quiet periods and low bag intensities (Figure 3, E and F), with the switch occurring between mApple intensity values of 10.1 and 14.8. This spatiotemporal analysis indicates that SPV-1 spatially regulates calcium activity in the spermatheca by keeping calcium low in the bag cells and sp-ut valve for a period of time after oocyte entry.

### Increasing RHO-1 activity recapitulates transit timing of the *spv-1* mutant, but not calcium signaling

Because SPV-1 acts through RHO-1 to regulate contractility of the spermatheca (Tan and Zaidel-Bar, 2015), we speculated that increasing RHO-1 activity might alter calcium signaling in a manner similar to the loss of SPV-1. To test this idea, we obtained nematodes expressing constitutively active RHO-1(G14V) under the control of a heat shock promoter (McMullan and Nurrish, 2011), crossed them with GCaMP expressing animals, and optimized the heat shock protocol to enable the capture of embryo transit movies (Figure 4, A and B). As expected, increasing RHO-1 activity results in shorter dwell times (Figure 4, A and C). However, increasing RHO-1 activity did not phenocopy *spv-1* mutant calcium signaling, showing slow rising times and low fractions over half max (Figure 4, A, B and C), and long sp-ut quiet periods and low bag intensities (Figure 4, D and E). In all calcium signaling metrics, increased RHO-1 activity did not differ significantly from the non-heat shocked control or from the wildtype population. These results suggest that SPV-1 is likely not working through RHO-1 to regulate calcium signaling in the spermatheca.

**Figure 4.**
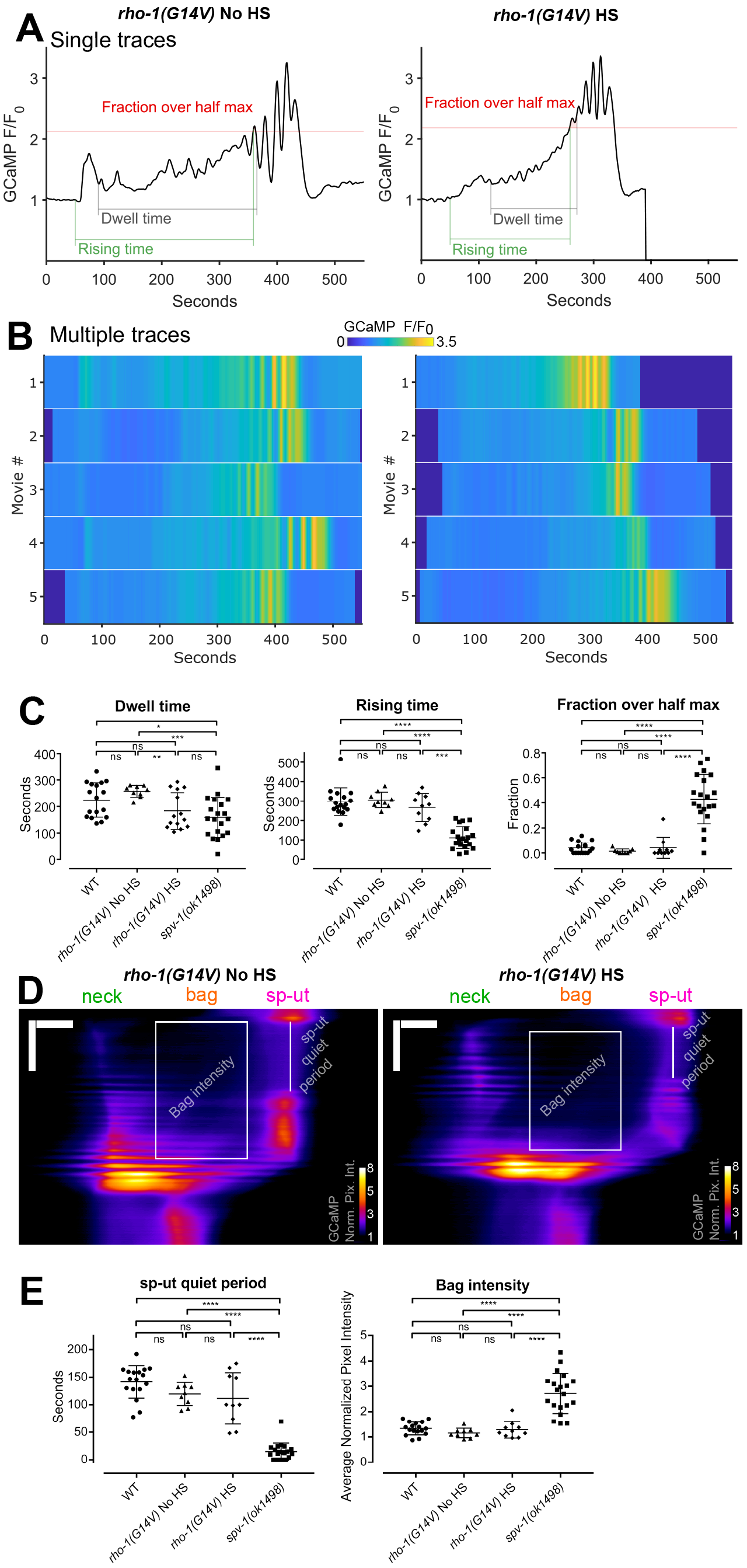
Increasing RHO-1 activity alters spermathecal contractility but does not recapitulate *spv-1(ok1498)* mutant calcium signaling. (A) Representative time series from embryo transits with metrics annotated. (B) Heat maps showing time series from multiple embryo transits. The time series in panel A corresponds to the first row of the heat map. (C) Quantification of time series metrics. (D) Representative kymograms with metrics annotated. Horizontal scale bars are 10 μm, vertical scale bars are 100 seconds. (E) Quantification of kymogram metrics. In panels C and E error bars display standard deviation, p-values were calculated using Welch’s ANOVA with Games-Howell multiple comparison: ns, p ≥ 0.05, *p < 0.05, **p < 0.01, ***p <0.001, ****p < 0.0001. WT and *spv-1(ok1498)* data are duplicated from figures 1 and 3.

### SPV-1 regulates calcium signaling via its GAP domain

A functional RhoGAP domain is required for SPV-1 to regulate spermathecal contractility and embryo transit timing (Tan and Zaidel-Bar, 2015). To explore if this activity is also needed for SPV-1 to regulate calcium signaling, we generated transgenic nematodes expressing labeled SPV-1 with a nonfunctional RhoGAP domain, SPV-1(R635K)::mApple, and acquired two-color movies of embryo transits. *spv-1(R635K)::mApple* was expressed at average mApple fluorescence intensities from 11 to 24, coinciding with mApple fluorescence levels at and above the threshold that induces trapping with RhoGAP functional SPV-1::mApple. Transits in *spv-1(ok1498); spv-1(R635K)::mApple* were similar to those seen in the *spv-1* mutant, with short rising times, high fractions over half max, short sp-ut quiet periods, and high bag intensities (Figure 5, A, B, C and D). These data suggest that the GAP activity of SPV-1 is required to modulate calcium signaling, and indicate that the downstream target of SPV-1 is likely to be a GTPase.

**Figure 5.**
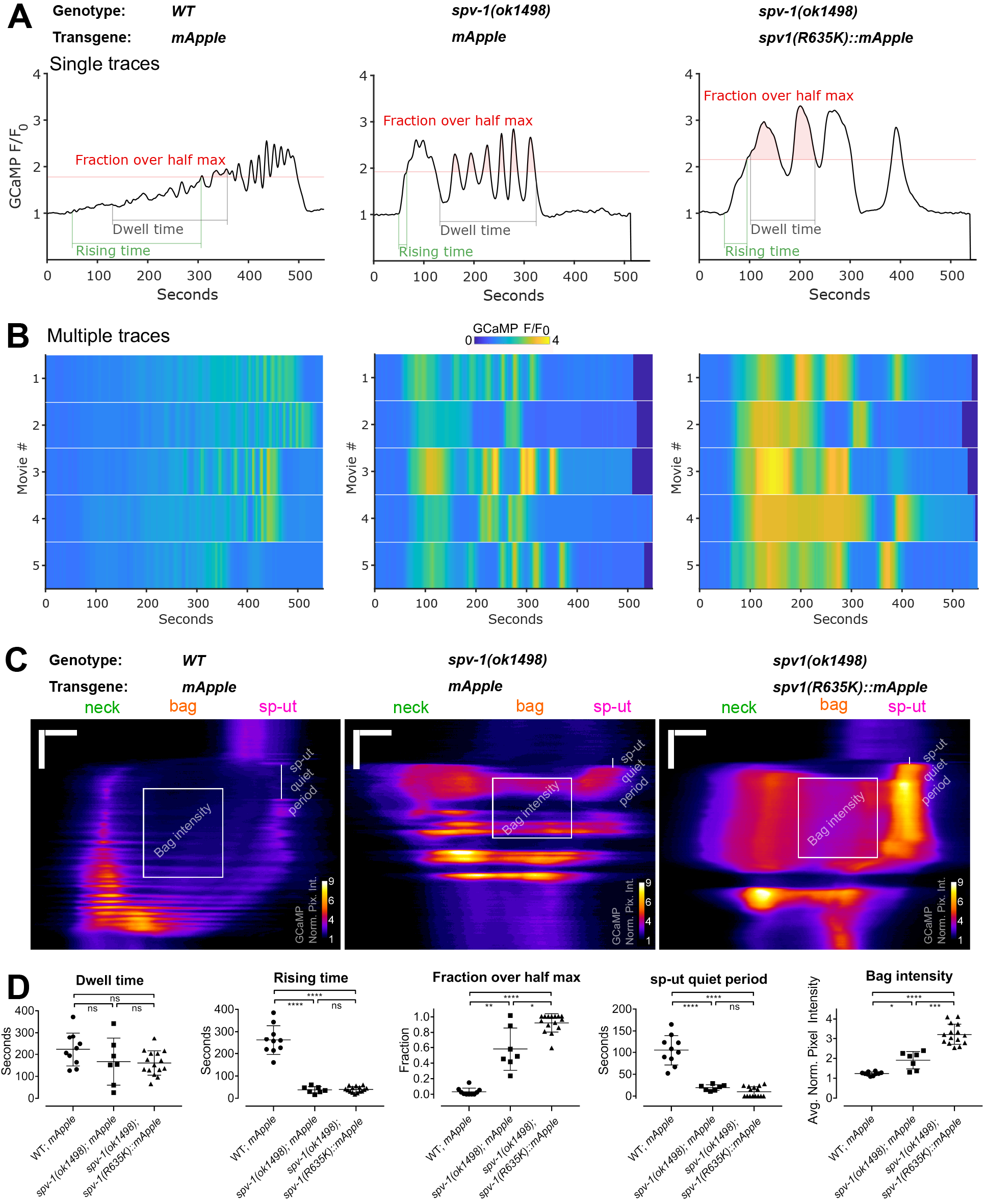
SPV-1 regulates calcium signaling through its GAP domain. (A) Representative time series from embryo transits with metrics annotated. (B) Heat maps showing time series from multiple embryo transits. Time series in panel A correspond to the first rows of the heat maps. (C) Representative kymograms with metrics annotated. Horizontal scale bars are 10 μm, vertical scale bars are 100 seconds. (D) Quantification of metrics. Error bars display standard deviation, p-values were calculated using Welch’s ANOVA with Games-Howell multiple comparison: ns, p ≥ 0.05, *p < 0.05, **p < 0.01, ***p <0.001, ****p < 0.0001. *WT; mApple* and *spv-1(ok1498); mApple* data are duplicated from figures 2 and 3.

### SPV-1 has GAP activity toward Cdc42 and partially co-localizes with CDC-42

In addition to regulating RHO-1, *in vitro* RhoGAP activity assays indicate SPV-1 has significant GAP activity toward another Rho family GTPase, Cdc42/CDC-42 (Figure 6A; (Ouellette et al., 2015)). To determine if CDC-42 is present in the cells of the spermatheca, we obtained and imaged nematodes expressing a CDC-42::GFP fusion (Neukomm et al., 2014) and found CDC-42 expressed throughout the spermatheca at the apical and basal membranes (Figure 6, B and C). To investigate the spatial relationship between CDC-42 and SPV-1, we crossed *spv-1::mApple* into the *cdc-42::GFP* animals and acquired two-color movies of embryo transits. We found that SPV-1::mApple partially co-localizes with CDC-42::GFP at the cell membranes (Figure 6, D and E). These data suggest that SPV-1 may regulate CDC-42 in spermathecal cells.

**Figure 6.**
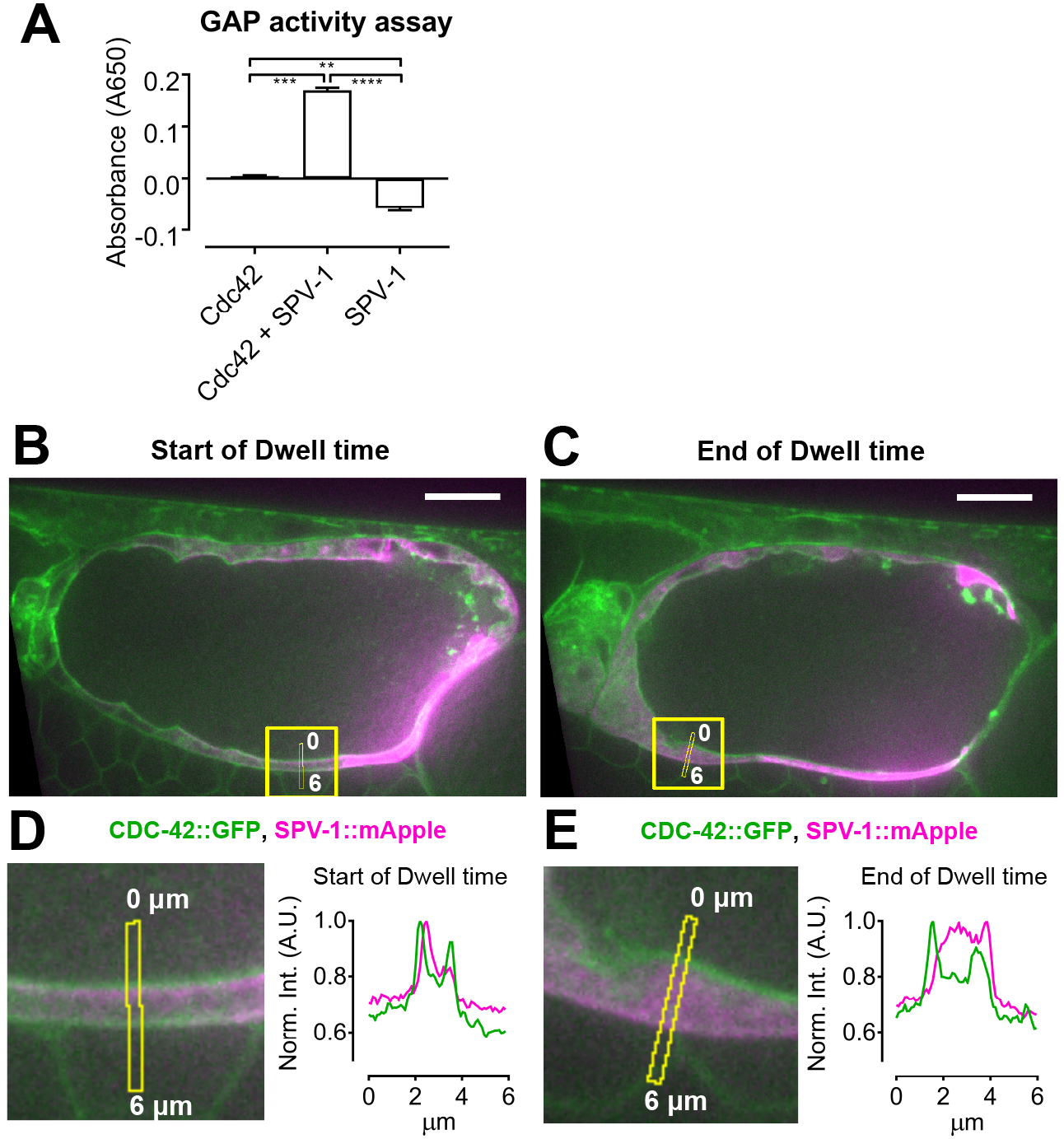
SPV-1 exhibits GAP activity toward Cdc42 and partially co-localizes with CDC-42 at spermathecal cell membranes.(A) In vitro assay measuring activity of the SPV-1 RhoGAP domain toward recombinant mammalian Cdc42. Error bars display S.E.M, p-values were calculated using Welch’s ANOVA with Games-Howell multiple comparison: **p < 0.01, ***p <0.001, ****p < 0.0001. (B, C) The first and last frames of the dwell time from a representative movie, with the quantified region, capturing the same cell, annotated. Scale bars are 10 μm. (D, E Left) Digitally zoomed view of the quantified region, with the line scans annotated. (D, E Right) Average fluorescence intensity along the line scans, with CDC-42::GFP in green and SPV-1::mApple in magenta.

### Increasing CDC-42 activity recapitulates many aspects of *spv-1* mutant calcium signaling

To determine if increasing CDC-42 activity alters calcium signaling similarly to the loss of SPV-1, we generated nematodes expressing a constitutively active CDC-42(Q61L) (Aceto et al., 2006; Ziman et al., 1991) under the control of a heat shock promoter, crossed this line with our GCaMP sensor line, optimized the heat shock protocol for CDC-42, and acquired embryo transit movies. Increasing CDC-42 activity does not significantly alter dwell times, although it does produce *spv-1* mutant-like calcium signaling with faster rising times, increased fractions over half max, shorter sp-ut quiet periods, and higher bag intensities (Figure 7, A, B, C, D and E). These data suggest SPV-1 likely acts through CDC-42 to regulate calcium signaling in the spermatheca during embryo transits.

**Figure 7.**
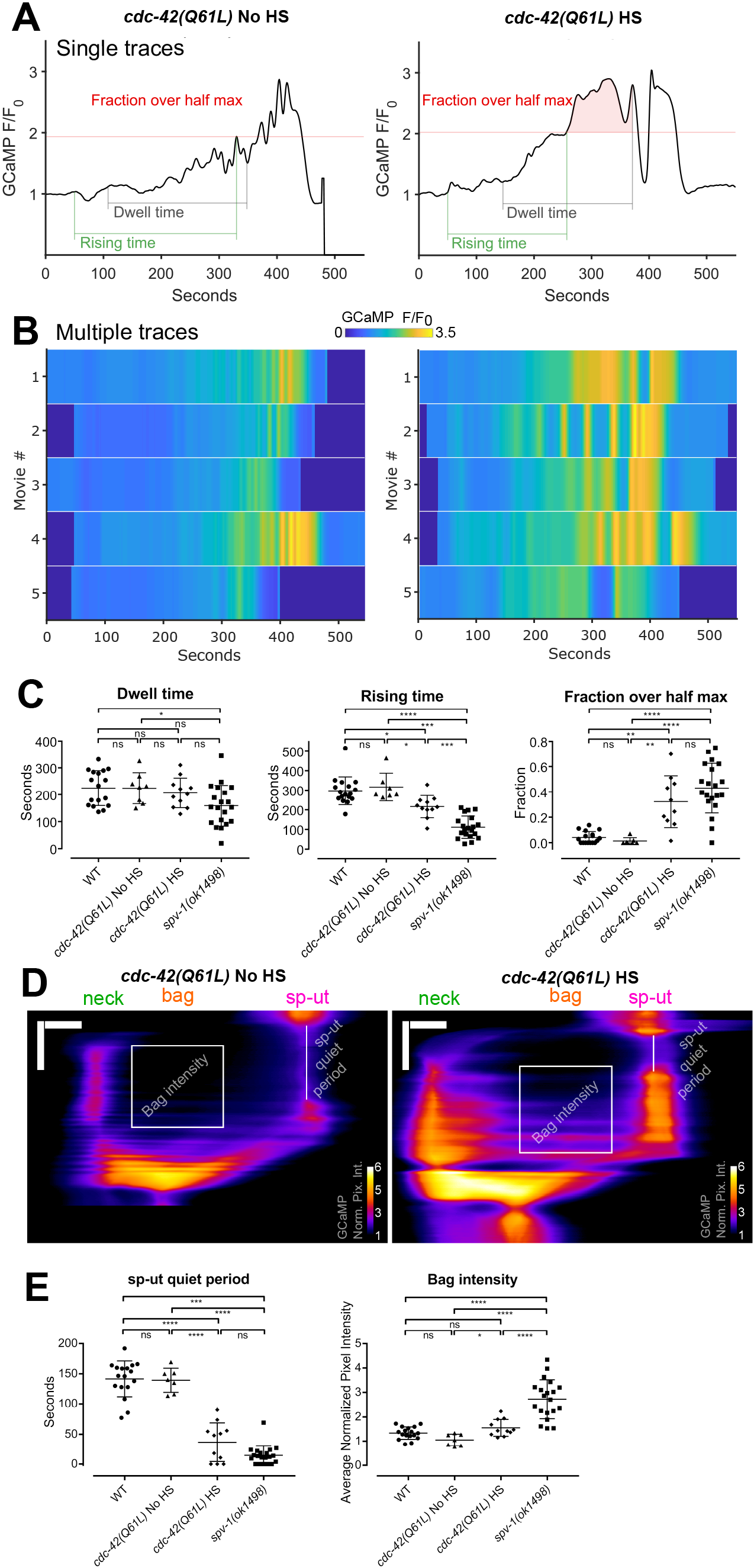
Increasing CDC-42 activity alters spermathecal calcium signaling. (A) Representative time series from embryo transits with metrics annotated. (B) Heat maps showing time series from multiple embryo transits. Time series in panel A corresponds to the first row of the heat map. (C) Quantification of time series metrics. (D) Representative kymograms with metrics annotated. Horizontal scale bars are 10 μm, vertical scale bars are 100 seconds. (E) Quantification of kymogram metrics. In panels C and E error bars display standard deviation, p-values were calculated using Welch’s ANOVA with Games-Howell multiple comparison: ns, p ≥ 0.05, *p < 0.05, **p < 0.01, ***p <0.001, ****p < 0.0001. WT and *spv-1(ok1498)* data are duplicated from figures 1 and 3.

### Decreasing CDC-42 using RNAi does not alter calcium signaling

To further explore CDC-42′s role in calcium signaling we generated nematodes coexpressing a red calcium sensor, R-GECO, and CDC-42 labeled with GFP. We used RNAi against *cdc-42* to deplete CDC-42 levels and recorded embryo transit movies with R-GECO to monitor calcium signaling. Decreasing CDC-42 levels via RNAi does not alter calcium time series in the *spv-1* mutant background (Figure 8, A and B), or in the wildtype background (data not shown). Dwell times and global calcium signaling metrics are unchanged by *cdc-42(RNAi)* in both wildtype and mutant backgrounds (Figure 8C). Spatiotemporal patterns of calcium signaling are also unchanged by *cdc-42(RNAi)*, indicated by kymograms (Figure 8D) and spatiotemporal calcium signaling metrics (Figure 8E). Measurements of CDC-42::GFP fluorescence intensity suggests that *cdc-42(RNAi)* treatments are significantly depleting CDC-42 levels (Figure 8F), however residual levels of CDC-42 could be sufficient to support the observed calcium dynamics. These results suggest that SPV-1 may regulate calcium signaling through another as yet unidentified effector.

**Figure 8.**
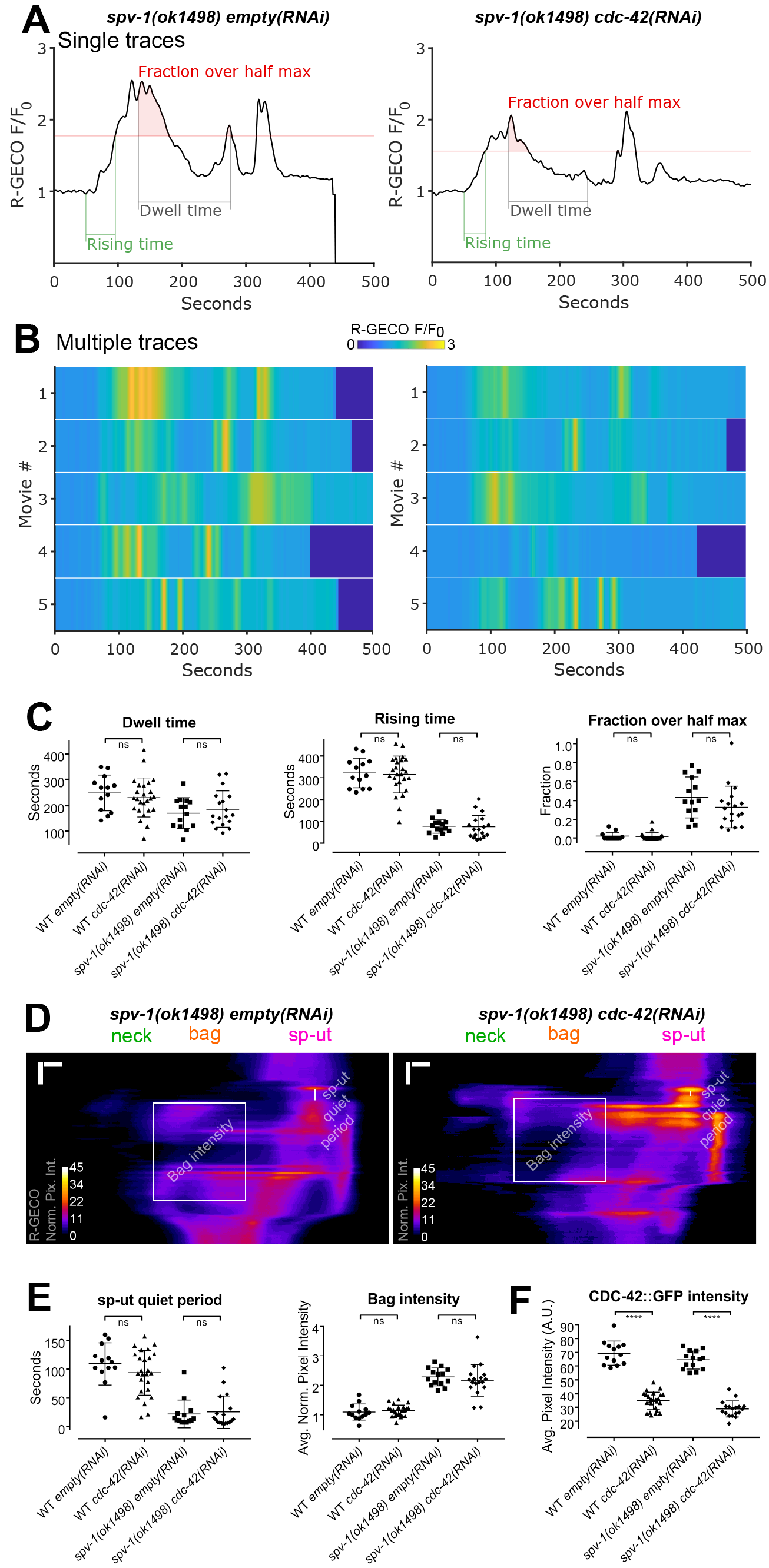
*cdc-42(RNAi)* does not alter calcium signaling. (A) Representative time series from embryo transits with metrics annotated. (B) Heat maps showing time series from multiple embryo transits. The time series in panel A corresponds to the first row of the heat map. (C) Quantification of time series metrics. (D) Representative kymograms with metrics annotated. Horizontal scale bars are 5 μm, vertical scale bars are 50 seconds. (E) Quantification of kymogram metrics. (F) Quantification of CDC-42::GFP intensity. In panels C, E and F error bars display standard deviation, p-values were calculated using Welch’s t-test: ns, p ≥ 0.05, ****p < 0.0001.

## Discussion

In this work, we show that the RhoGAP SPV-1 is a major regulator of calcium signaling in the *C. elegans* spermatheca during embryo transits. SPV-1 controls spatiotemporal aspects of calcium signaling by keeping calcium low in the spermathecal bag cells and sp-ut valve for a period of time after oocyte entry. This controlled calcium signaling likely contributes to the spatiotemporal patterns of contractility present in wildtype function of the tissue, where contractility must be high in the distal neck to keep the embryo in the spermatheca, and yet must also stay low in the central bag cells so that the egg forms with the correct shape. Misregulated calcium signaling, in addition to increased active RHO-1 levels, likely contributes to the misshapen embryos and decreased brood sizes seen in the *spv-1(ok1498)* mutant (Tan and Zaidel-Bar, 2015).

Spermathecal tissue function and calcium signaling exhibit a threshold response to SPV-1::mApple level, suggesting levels of SPV-1 protein must be maintained within a narrow range for wildtype embryo transits to occur. Increases in SPV-1 above wildtype levels lead to over-dampening of the signaling pathways regulating contractility, resulting in a tissue that cannot produce productive contractions. Decreases in SPV-1 below wildtype levels lead to insufficient inhibition of the signaling pathways that stimulate contractility, making the tissue hypercontractile (Figure 9, A and B). A threshold response also suggests that feedback mechanisms are involved in controlling spermathecal tissue function and calcium signaling. The molecular mechanisms tightly regulating the expression and activity levels of SPV-1, and characterization of the feedback control system that governs the spermatheca, are exciting areas for future investigation.

**Figure 9.**
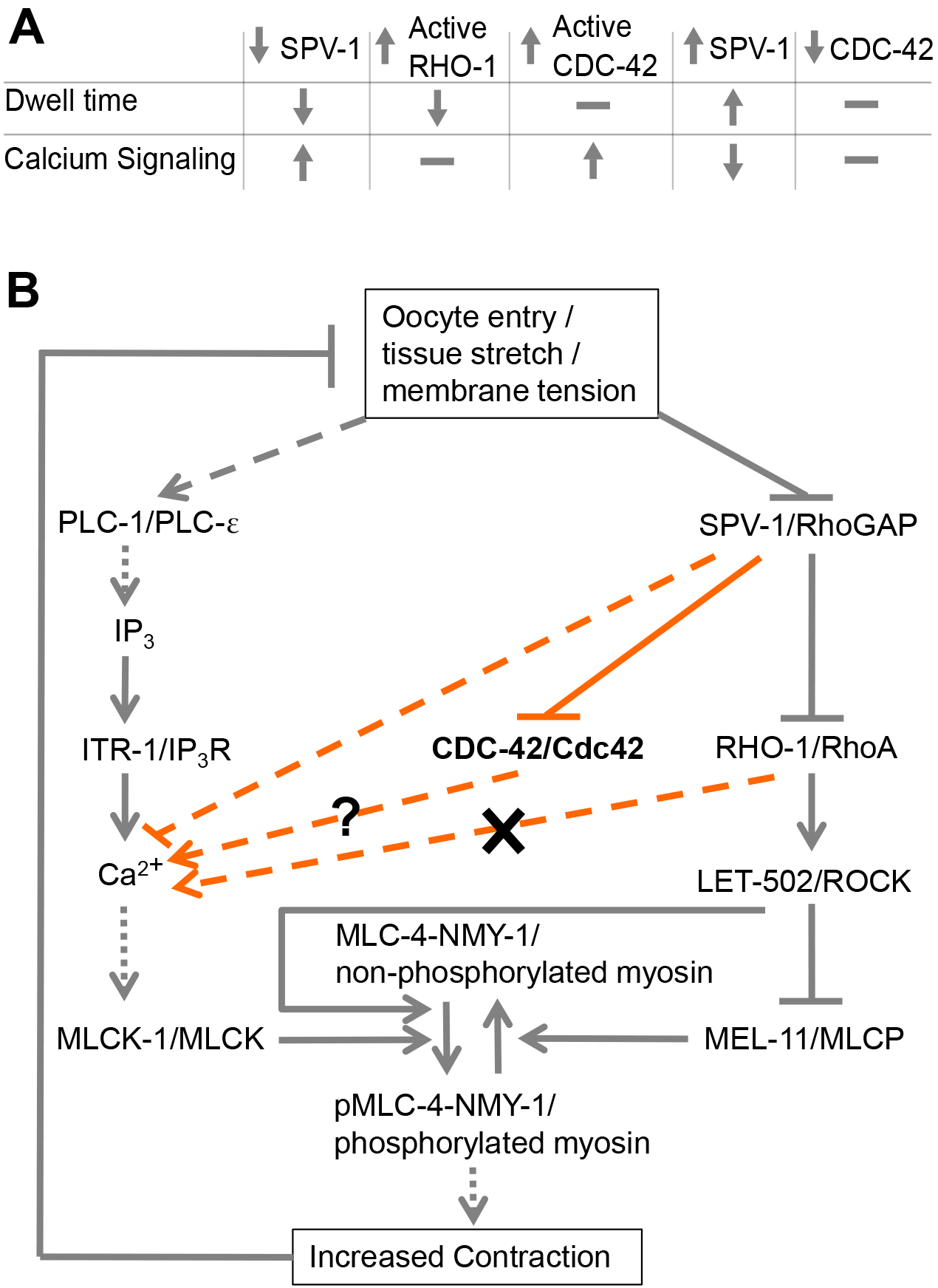
SPV-1 regulates spermathecal contractility via calcium and Rho-ROCK signaling. (A) Summary table of findings. SPV-1 regulates both the Rho-ROCK and calcium signaling pathways which together are required for activation of myosin and tissue contractility. Increasing RHO-1 activity leads to faster transits (decreased dwell times) similar to decreased SPV-1, but does not alter calcium signaling. Increasing active CDC-42 does not alter dwell times but does alter calcium signaling similar to decreased SPV-1. Increasing SPV-1 increases dwell times, often resulting in transit failures and embryo trapping, and decreases calcium signaling activity and magnitude. Decreasing CDC-42 using RNAi does not alter dwell time or calcium signaling. (B) Proposed model of the network regulating actomyosin contractility in the spermatheca. Grey lines indicate previously known interactions, orange lines indicate interactions found in this study. Solid lines indicate direct interactions, dotted lines indicate resultant interactions with known intermediates not shown, and dashed lines indicate unknown intermediates.

Given previous results suggesting SPV-1 acts through RHO-1 to regulate contractility, it was a surprise to find that constitutively active RHO-1 did not recapitulate the *spv-1(ok1498)* calcium signaling phenotypes. Although animals expressing constitutively active RHO-1 did exhibit shorter dwell times, they produced fairly wildtype calcium signaling. RHO-1 is therefore probably working predominantly through LET-502/ROCK and myosin activation, and not in the regulation of calcium signaling. Increased phosphorylation of myosin through the Rho-ROCK pathway would lead to calcium sensitization of myosin, a common phenomenon in smooth muscle and non-muscle systems in which contraction is initiated at lower than normal calcium levels (Somlyo and Somlyo, 2003).

SPV-1 requires a functional GAP domain to regulate calcium during embryo transits, suggesting a GTPase is modulating calcium signaling in the spermatheca. Animals expressing constitutively active CDC-42 in the spermatheca recapitulated many aspects of the *spv-1* mutant calcium signaling phenotypes, although the dwell times were unchanged, indicating that premature and elevated cytosolic calcium alone cannot generate the hyperconstriction of the spermatheca and rapid embryo transits observed in *spv-1(ok1498)* animals.

Animals expressing constitutively active CDC-42 show bag intensities which are significantly different from the non-heat shocked control, but not from the wildtype population. This could be due to the expression of constitutively active CDC-42 throughout the tissue instead of active CDC-42 following spatiotemporal patterns, or it could be that the heat shock driven expression of constitutively active CDC-42 was not high enough to see the full response. Like the bag intensities, the rising times are significantly different from the *spv-1* mutant population, and this uneven recapitulation of the *spv-1* mutant calcium signaling phenotype may hint that there is another protein interacting with SPV-1 to regulate calcium signaling. However, the robust response in the fractions over half max and the sp-ut quiet periods indicate that increased CDC-42 activity can partially recapitulate the *spv-1* mutant calcium signaling phenotype.

If SPV-1 works through the inactivation of CDC-42 to regulate calcium release, depletion of CDC-42 might be expected to suppress the *spv-1* mutant calcium dynamics. However, depleting CDC-42 using RNAi did not result in altered calcium signaling in the *spv-1* mutant or wildtype backgrounds, despite evidence that the treatment is effective at depleting CDC-42. CDC-42 is important for larval development, including proper formation of the gonad (Norman et al., 2005), and depleting it completely results in morphogenesis defects that preclude ovulation. Due to this our RNAi treatments were started at later larval stages, and were not complete knockdowns. Perhaps residual CDC-42 is sufficient to properly regulate calcium signaling. Another possibility is that SPV-1 regulates calcium through another unknown effector. Our results demonstrate the GAP activity of SPV-1 is required for proper spatiotemporal regulation of calcium signaling, suggesting that a GTPase is a target. Of the known Rho GTPases in *C. elegans* (CED-10, RAC-2, MIG-2, CDC-42, RHO-1, and CRP-1), SPV-1 has been shown to only have significant GAP activity toward CDC-42 and RHO-1 (Ouellette et al., 2016). Identification of additional SPV-1 effectors will be an important avenue for future studies.

In other systems, CDC-42/Cdc42 is known to regulate actin cytoskeletal rearrangements and cell migration as well as protein kinase cascades that can alter transcription of downstream targets (Johnson, 1999; Takai et al., 2001). However, few studies link Cdc42 to the activation of calcium signaling. Perhaps the best characterized example is found in mast cells, a critical cell in the immune system during inflammatory reactions. Calcium signaling, triggered by antigen binding to IgE and Fc_ε_RI receptors, controls mast cell exocytosis and degranulation, which releases histamines and other substances. In RBL-2H3mast cells, dominant active mutants of Cdc42 lead to elevated levels of antigen stimulated IP_3_ and cytosolic calcium (Hong-Geller and Cerione, 2000). Further studies have shown that Cdc42 can participate in the regulation of stimulated PIP_2_ synthesis, the substrate of PLC isozymes, in these cells (Wilkes et al., 2014). This suggests CDC-42 activity in the spermatheca may be controlling calcium activity through the regulation of PIP_2_ turnover and/or the production of IP_3_. In addition, *in vitro* studies have shown that human Cdc42 can stimulate purified PLC-β (Illenberger et al., 1998) and interact with PLC-γ1 (Hong-Geller and Cerione, 2000). It is possible that CDC-42 is affecting the activation of PLC-1/ε or the availability of substrate in the spermathecal cells to control IP_3_ production and calcium release during embryo transits.

SPV-1 has three human orthologs (Tan and Zaidel-Bar, 2015): ARHGAP29/PARG1 (Myagmar et al., 2005; Saras et al., 1997), HMHA1/ARHGAP45 (Amado-Azevedo et al., 2018; de Kreuk et al., 2013) and GMIP/ARHGAP46 (Aresta et al., 2002). Recently, loss of HMHA1/ARHGAP45 in endothelial cells was found to increase wound healing and cell migration rates, suggesting altered actomyosin contractility, as well as increasing the cellular response to shear stress, suggesting altered mechanotransduction (Amado-Azevedo et al., 2018). Similar patterns of altered mechanotransduction and contractility are present in the spermatheca when SPV-1 is lost. In mice, ARHGAP29/PARG1 and a binding partner Rasip1 were found to regulate nonmuscle myosin II via RhoA and Cdc42 during blood vessel development, with Rasip1 first activating Cdc42 to break cell-cell adhesions during lumen formation and Rasip1 then activating ARHGAP29/PARG1, inhibiting RhoA and keeping contractility low so the lumen stays open and the blood vessel can expand as the animal grows (Barry et al., 2016). This behavior of ARHGAP29/PARG1, regulating cellular contractility directly through RhoA and also through Cdc42 in conjunction with a binding partner, further suggests other proteins may interact with SPV-1 to modulate calcium signaling in the spermatheca.

Our results indicate SPV-1 regulates calcium signaling in addition to Rho-ROCK signaling, providing insights into the regulation of calcium signaling, actomyosin contractility, and tissue function in the *C. elegans* spermatheca. From this evidence it appears that not only is there interaction between the two central pathways that regulate actomyosin contractility, but a single protein can fine-tune both pathways to generate consistent and robust tissue function. Given conservations of sequence and function, it will be exciting to determine whether SPV-1 orthologs play similar roles in other biological contexts.

## Acknowledgements

We thank Ismar Kovacevic, Pei Yi Tan, Anand Asthagiri, Javier Apfeld, and members of the Cram and Apfeld labs for helpful feedback and discussions. This work was supported by a grant from the National Institutes of Health/National Institute of General Medical Sciences (GM110268) to E.J.C., a grant from the Israel Science Foundation (grant No. 1293/17) to R.Z-B., a National Science Foundation/Molecular and Cellular Biosciences - U.S. Israel Binational Science Foundation award (1816640) to E.J.C. and R.Z-B., and a National Science Foundation East Asia and Pacific Summer Institutes fellowship (1414889) to J.B. Some *C. elegans* strains were provided by the *Caenorhabditis* Genetics Center, which is funded by the National Institutes of Health Office of Research Infrastructure Programs (P40 OD010440).

## Methods

### *C. elegans* Strains and Culture

Nematodes were grown on nematode growth media (NGM) (0.107 M NaCl, 0.25%wt/vol Peptone (Fisher), 1.7% wt/vol BD Bacto-Agar (Fisher), 0.5% Nystatin (Sigma), 0.1 mM CaCl_2_, 0.1 mM MgSO_4_, 0.5% wt/vol cholesterol, 2.5 mM KPO_4_), and seeded with *Escheria coli OP50* using standard techniques (Myers et al., 1996). Nematodes were cultured at 20°C unless specified otherwise. All lines expressing GCaMP3 were crossed into UN1108 as described in Kovacevic et al. (2013) or UN1417, a separate integration event of the same construct, *fln-1p::GCaMP3*. GCaMP signal in UN1108 and UN1417 does not differ significantly in any of the described metrics. *spv-1(ok1498)* animals were crossed with UN1417 animals to generate the line UN1416 (*spv-1(ok1498); fln-1p::GCaMP3*). Table 1 provides a full list of *C. elegans* strains used in this study.

**Table 1.**
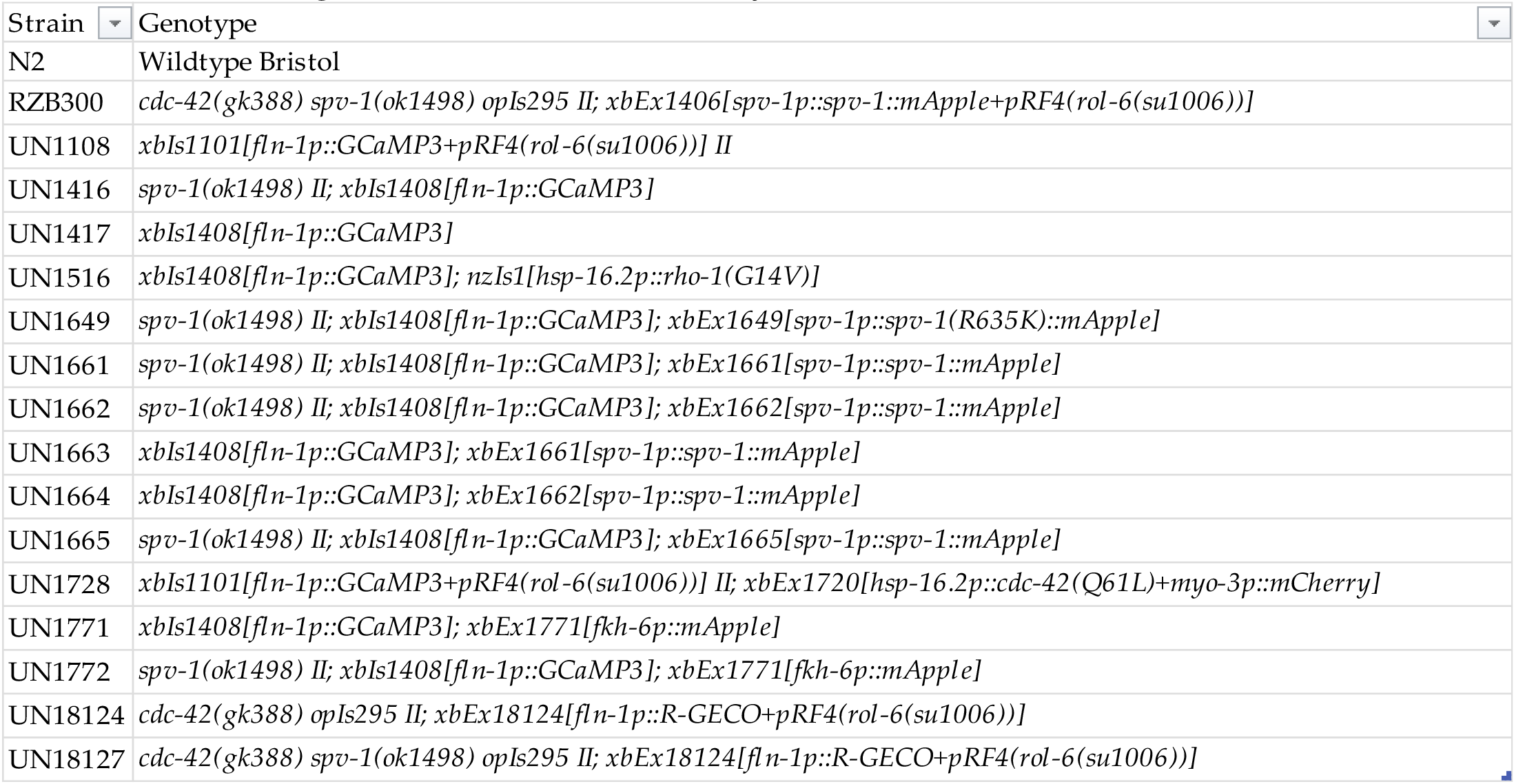
List of *C. elegans* strains used in this study.

#### Construction of spv-1::mApple and SPV-1(R635K)::mApple transgenic animals

*GFP* and the *spv-1* 3′UTR were removed from plasmid pPY1 (*spv-1p::spv-1::GFP::spv-1 3′UTR*, Tan and Zaidel-Bar 2015) and replaced with *tdTomato* and the *fln-1* 3′UTR from pUN284 (*fln-1p::inx-12::tdTomato::fln-1 3′UTR*) using restriction digest cloning with the enzymes KpnI and EagI. *tdTomato* was removed from this plasmid using the enzymes KpnI and NotI, and replaced with *mApple* amplified from Addgene plasmid 27698 using primers that introduced KpnI and NotI restriction sites to the ends of *mApple*, generating pUN362. *spv-1p::spv-1::mApple::fln-1 3′UTR* was then transferred from pUN362 into pUN359, a plasmid modified from pCFJ151 (Frøkjær-jensen et al., 2008) to facilitate CRISPR insertion at the *ttTi5605* transposon site, using restriction digest cloning with enzymes PmeI and SpeI, resulting in pUN427. pUN595 (*spv-1p::SPV-1*(*R635K*)::*mApple*::*fln-1 3′UTR*) was generated using around the horn PCR site-directed mutagenesis on pUN427 with primers designed to introduce the two nucleotide R635K amino acid mutation as well as a single nucleotide silent mutation 30 bp downstream to generate a BglII restriction site to facilitate tracking the R635K mutation. Transgenic animals were created by microinjecting N2 nematodes with a DNA solution containing 50 ng/μL of Cas9 plasmid (Chen et al., 2008; Tzur et al., 2013), 30 ng/μL each of 2 different CRISPR guide plasmids, pUN357 and pUN358, and 45 ng/μL of pUN427 or pUN595. Progeny displaying red fluorescence in the spermatheca were isolated and screened for CRISPR integration. After multiple failed attempts at CRISPR integration, lines expressing the transgenes as extrachromosomal arrays were established and used for experiments. Three *spv-1::mApple* lines and two *SPV-1(R635K)::mApple* lines were isolated from independent microinjections. These lines were crossed with UN1416 nematodes to generate animals with dual-labeled spermathecae in the wildtype and *spv-1* mutant backgrounds.

#### Construction of fkh-6p::mApple transgenic animals

*GFP* was removed from pUN106 (*fkh-6p::GFP::unc-54 3′UTR*) and replaced with *mApple* from pUN427 using restriction digest cloning with enzymes KpnI and EcoRI to create pUN822. Transgenic animals were created by microinjecting N2 nematodes with a DNA solution containing 45 ng/μL of pUN822 and 100 ng/μL of pRF4 (*rol-6* injection marker). Progeny displaying red fluorescence in the spermatheca were isolated and lines expressing the transgene as an extrachromosomal array were established. These lines were crossed with UN1416 nematodes to generate animals with dual-labeled spermathecae in the wildtype and *spv-1* mutant backgrounds.

#### Construction of constitutively active CDC-42 (hsp16.2p::CDC-42(Q61L)::unc-54 3′UTR) animals

*CDC-42(Q61L)* was amplified from pJK6 (a gift from Ahna Skop, made originally by John White) using primers engineered with EcoRI 5′ extensions and ligated into pUN597 (*hsp16.2::unc-54 3′UTR*) between the promoter and 3′UTR to create pUN624. Transgenic animals were created by microinjecting a DNA solution containing 20 ng/μl pUN624 and 25 ng/μl *myo-3:mCherry* as a co-injection marker into N2 animals. Animals exhibiting *mCherry* expression in the body wall muscle were segregated to establish the transgenic line UN1720. UN1720 was crossed into UN1108 (*fln-1p::GCaMP3*) creating the transgenic line UN1728 for calcium studies.

#### Construction of cdc-42::GFP: spv-1(ok1498): spv-1::mApple animals

The strain WS5018 (*cdc-42(gk388);opIs295 II (cdc-42::gfp*)) (Neukomm et al., 2014) was obtained from the *Caenorhabditis* Genetics Center (CGC) and crossed with a *spv-1(ok1498); spv-1::mApple* line generated above, creating strain RZB300.

#### Construction of cdc-42::GFP: R-GECO animals

WS5018 animals were injected with a DNA solution containing 50 ng/μl of pUN526 (*fln-1p::R-GECO::fln-1 3′UTR*) and 40 ng/μl of pRF4 (*rol-6* injection marker). Progeny displaying red fluorescence in the spermatheca were isolated and a line, UN18124, expressing the transgene as an extrachromosomal array was established. RZB300 animals lacking red fluorescence were crossed with UN18124 animals to generate the strain UN18127 with *cdc-42::GFP and R-GECO* in the *spv-1(ok1498)* background.

### Heat shock protocols for constitutively active *rho-1(G14V)* and *cdc-42(Q61L)*

*UN1516 (RHO-1(G14V); fln-1p::GCaMP3*) animals were synchronized using an ′egg prep′ where embryos are released from young gravid adults using an alkaline hypochlorite solution (Hope, 1999). Clean embryos were plated on *OP50* seeded NGM and cultured at 20°C until young adulthood (70-72 hours post ′egg prep′). To induce expression of constitutively active *RHO-1(G14V)*, animals were moved from 20°C to 33°C for 30 minutes, and then left to recover at 20°C for 1 hour before imaging.

*UN1728 (CDC-42(Q61L); fln-1p::GCaMP3)* animals were synchronized as described above. Young adults expressing *myo-3::mCherry* were segregated and placed at 33°C for 2 hours to induce the expression of constitutively active *CDC-42(Q61L)*. Animals were left to recover at 20°C for 1 hour before imaging.

### RNA interference

The RNAi protocol was performed essentially as described in (Timmons and Fire, 1998). HT115(DE3) bacteria transformed with the dsRNA construct of interest were grown overnight in Luria Broth (LB) supplemented with 40 mg/ml ampicillin. The following day 300 μl of the cultured bacteria was seeded on NGM/RNAi plates supplemented with 25 μg/ml carbenicillin and disopropylthio-β-galactoside (IPTG) and left for 24-72 hours to induce dsRNA expression. Diluted *spv-1* RNAi was conducted by mixing overnight cultures of *spv-1* and empty RNAi bacteria in volume ratios of 1:1, 1:3, 1:9, or 1:19 for seeding NGM/RNAi plates. Embryos were collected using an ′egg prep′ as described above. Clean embryos were transferred to the seeded NGM/RNAi plates and incubated at 20°C for 65-70 hours before imaging. For *cdc-42* RNAi clean embryos were deposited on *OP50* NGM plates and incubated at 20°C for 36-48 hours, followed by transfer to *cdc-42* or empty RNAi plates and incubation at 20°C for another 24-36 hours before imaging.

### Image Acquisition

For all GCaMP and R-GECO studies, animals were immobilized with 0.05-micron polystyrene beads (Polysciences Inc., Warrington, PA, USA), mounted on slides with 5-10% agarose pads (Kim et al., 2013), and imaged using a 60×, 1.40 NA oil-immersion objective on a Nikon Eclipse 80i microscope equipped with a SPOT RT3 CCD camera (Diagnostic instruments; Sterling Heights, MI, USA) controlled by SPOT Advanced imaging software, version 5.0, with Peripheral Devices and Quantitative Imaging modules. Images were acquired at 1600×1200 pixels, using the full camera chip, and saved as 8-bit tiff files. Fluorescence excitation was provided by a Nikon Intensilight C-HGFI130W mercury lamp and shuttered with a Lambda 10-B SmartShutter (Sutter Instruments, Novato CA, USA), also controlled through the SPOT software. Single channel GCaMP time lapse movies were acquired using a GFP filter set (470/40× 495bs 525/50m) (Chroma Technologies, Bellows Falls VT, USA) at 1 frame per second, with an exposure time of 75 ms, gain of 8, and neutral density of 16. Two channel GCaMP/mApple and GFP/R-GECO time lapse movies were acquired using an EGFP/mCherry filter set, with excitation filters 470/40× and 572/35× housed in the SmartShutter filterwheel to enable rapid switching, and emissions passing through a single filter cube containing beamsplitter 59022bs and emission filter 59022m (520/20m and 640/40m) (Chroma Technologies). For GCaMP/mApple movies one GCaMP frame and one mApple frame were acquired every 2 seconds, with exposure time of 40 ms and gain of 16 for the GCaMP channel, exposure time of 75 ms and gain of 32 for the mApple channel, and neutral density of 32 for both channels. For GFP/R-GECO movies one GFP frame and one R-GECO frame were acquired every 2 seconds, with exposure time of 50 ms, gain of 16, and neutral density of 16 for both channels.

For CDC-42::GFP/SPV-1::mApple studies, animals were mounted with 2 μl of M9 buffer on slides with 10% agarose pads, and imaged using a 100× 1.4 NA oil-immersion objective on a Nikon Ti confocal microscope equipped with a spinning-disk head (CSU-X1; Yokogawa, Tokyo, Japan) and Prime95b camera (Photometrics, Tucson, AZ) controlled using Metamorph software (Molecular devices, Sunnyvale, CA), and saved as 16-bit tiff files. Z-stacks were acquired with 1 μm spacing between slices, with Z-stacks acquired every 10 seconds. GFP was excited using the 481 nm laser and mApple was excited with the 561 nm laser, both at 35% laser power and 200 ms exposure time.

### Image Processing

Only successful embryo transits, meaning the embryo exited through the sp-ut valve, or whole embryos trapped in the spermatheca, were analyzed in this work. GCaMP and R-GECO movies were processed using a custom Fiji script. First, the movie was registered using the Fiji plugin StackReg (Thévenaz et al., 1998) to correct for any movement of the animal during image acquisition. The movie was then rotated to a standard orientation with the occupied spermatheca horizontal and the sp-ut valve on the right side of the movie. Finally, the movie was cropped to 800×400 pixels, annotated as processed, and saved as an 8-bit tiff file. For GCaMP/mApple and GFP/R-GECO movies the2 channels were registered separately and then recombined before rotation and cropping. The angle of rotation and positioning of the cropping box were determined by manual user input.

CDC-42::GFP/SPV-1::mApple movies were not processed beyond selecting a single Z-plane for analysis. Fiji was used to manually track and annotate the cells, and to manually draw and quantify line scans.

### Generation of GCaMP and R-GECO time series

GCaMP and R-GECO time series were generated by calculating the average pixel intensity for each frame of the processed movie. Normalized average pixel intensity time series were generated by normalizing the entire time series to the baseline value, F_0_, calculated as the average of the 30 frames prior to the start of oocyte entry (Kovacevic 2013).

### Extraction of metrics from time series

Custom Fiji and Matlab code was used to document and archive manually annotated time points, computationally identified metrics, and additional data for each movie. The global calcium signaling metrics rising time and fraction over half max were adapted from (Christo et al., 2015).

#### Manual annotation of time points, calculation of dwell time

Time points were determined by visual inspection. Four significant time points were annotated for each movie: (1) distal neck open, when the oocyte starts to enter the spermatheca, (2) distal neck close, when the embryo is completely enclosed, (3) sp-ut valve open, when the embryo starts to exit, and (4) sp-ut valve close, when the embryo completely exits the spermatheca. In ambiguous cases, three individuals scored each time point, and the average of the three values was used. Dwell time is calculated as the frame when the sp-ut valve opens minus the frame when the distal neck closes.

#### Calculation of max, half max and rising time

Each GCaMP and R-GECO time series was smoothed using a moving average filter, Matlab function ′filter′, with a window size of 5. The max value was determined using the built-in Matlab function, and the baseline value, previously used to normalize the time series, was recalled. The half max value was calculated by taking half the difference between the max and the baseline, and adding the baseline again. The first time point where the time series is above the half max value was recorded as the end of the rising time, with distal neck open as the start of the rising time. Rising time is calculated as first time point above half max minus distal neck open.

#### Calculation o f fraction over half max

For the dwell time of each time series, the number of time points above the half max value were counted, and this number was divided by the total dwell time to give the fraction over half max.

#### Calculation of mApple and CDC-42::GFP average pixel intensity

For GCaMP/mApple and CDC-42::GFP/R-GECO movies, mApple and CDC-42::GFP time series were generated by calculating the average pixel intensity of the mApple or GFP channel for each frame of the processed movie. The average of the 30 frames prior to oocyte entry, analogous to the GCaMP/R-GECO baseline, was recorded as the value for the mApple or CDC-42::GFP average pixel intensity. For presentation in Figure 2 the mApple time series was smoothed using a moving average filter with a window size of 50, and a constant 2 was added to every value of the time series to accommodate for the decreased intensity of the higher averaging. GCaMP/mApple movies were acquired in three batches over fifteen months, with a microscope deep cleaning and fluorescent lamp replacement between batches. SPV-1(R635K)::mApple average pixel intensity values were found to be consistent within batches, so the average SPV-1(R635K)::mApple average pixel intensity value for each batch was used to establish ratios to bring all the mApple values from the first two batches into agreement with the third batch.

#### Adjustment of metrics for trapped movies

Trapped dwell times were assigned a value of 575, slightly above the highest value in the dataset, for plotting purposes and fitting of the Hill function. These trapped dwell times were excluded from statistical analysis. Trapped fractions over half max were assigned a value of −0.10 for plotting purposes, and excluded from statistical analysis.

#### Fit to Hill_function and determination of SPV-1::mApple threshold

Using GraphPad Prism 7.04, dwell times were arranged in an XY table with mApple values as X and dwell times as Y, with all *spv-1(ok1498); spv-1::mApple* movies, i.e. trapped, untreated and RNAi, in a single column. Nonlinear regression curve fit was used, with [Agonist] vs. response – Variable slope (four parameters). The EC50 in the resulting table is presented here as the threshold.

### Generation of kymograms

For every frame of the processed movie the pixel intensities of each column were averaged to generate a single pixel representing that column. This operation was performed for each column of the frame, condensing the frame into a single pixel line. This operation was carried out for every frame of the movie, with the single pixel lines stacking to generate a 2D image that represents the spatiotemporal dynamics of the entire movie (Fig. 3 A, B, Movie 2). This was implemented in custom Fiji code using the commands Image>Stacks>Reslice followed by Image>Stacks>Z Project (Average Intensity).

### Extraction of spatiotemporal calcium signaling metrics from kymograms

The sp-ut quiet period was visually determined and indicated as a line drawn from the end of the first calcium transient to the start of the next rise in calcium level. The sp-ut quiet period was then calculated as the vertical distance between the endpoints of the line, measuring in seconds.

Before calculation of the bag intensity the kymogram was normalized by dividing the 32-bit kymogram image by the GCaMP or R-GECO time series baseline value, calculated above as the average of the 30 frames prior to the start of oocyte entry. A left bound at the distal neck and a right bound at the sp-ut valve were then identified. The bag intensity was calculated within a rectangle computationally selected to represent the lowest average calcium signal in a 25 μm region of the spermathecal bag during the dwell time. For trapped SPV-1::mApple kymograms, the average dwell time of the WT;mApple controls was used.

### In vitro RhoGAP activity assay

The RhoGAP activity assay was conducted as previously described (Tan and Zaidel-Bar, 2015).

### Statistical Analysis

Statistical tests were conducted at a significance level of 0.05. When 2 groups were compared, p-values were calculated using Welch′s t-test in GraphPad Prism. When more than 2 groups were compared, p-values were calculated using Welch′s ANOVA with Games-Howell multiple comparisons, in R (R Core Team, 2017), using the ′oneway′ function in the package ′userfriendlyscience′ (Peters, 2017).

## Movie legends

Movie 1. GCaMP embryo transit movies and time series display calcium signaling differences in *spv-1* mutants. Processed GCaMP embryo transit movies are shown in the bottom panels, while time series are simultaneously displayed in the top panels. A wildtype movie and time series are shown in the left column, while a *spv-1* mutant movie and time series are shown in the right column. Metric annotations appear in the time series as they occur in the movies. Both movies and time series were temporally aligned to put oocyte entry at 50 seconds.

Movie 2. Kymograms display the spatiotemporal differences in *spv-1* mutant calcium signaling. Processed GCaMP embryo transit movies, the same shown in Movie 1, are shown in the top panels, while kymograms are simultaneously displayed in the lower panels. A wildtype movie and kymogram are shown in the left column, while a *spv-1* mutant movie and kymogram are shown in the right column. Metrics appear on the kymograms as they occur in the movies. Both movies and kymograms were temporally aligned to put oocyte entry at 50 seconds.

## Abbreviations used

RhoGAP: GTPase activating protein toward Rho family small GTPases
ROCK: Rho-associated protein kinase
sp-ut: spermatheca uterine valve.

